# Sex-specific Associations of Gene Expression with Alzheimer’s Disease Neuropathology and Ante-mortem Cognitive Performance

**DOI:** 10.1101/2025.01.02.631098

**Authors:** Mabel Seto, Michelle Clifton, Melisa Lara Gomez, Gillian Coughlan, Katherine A. Gifford, Angela L. Jefferson, Philip L. De Jager, David A Bennett, Yanling Wang, Lisa L. Barnes, Julie A Schneider, Timothy J. Hohman, Rachel F. Buckley, Logan Dumitrescu

## Abstract

The biological mechanisms underlying women’s increased Alzheimer’s disease (AD) prevalence remain undefined. Previous case/control studies have identified sex-biased molecular pathways, but sex-specific relationships between gene expression and AD endophenotypes, particularly sex chromosomes, are underexplored. With bulk transcriptomic data across 3 brain regions from 767 decedents, we investigated sex-specific associations between gene expression and post-mortem β-amyloid and tau as well as antemortem longitudinal cognition. Of 23,118 significant gene associations, 10% were significant in one sex and not the other (sex-specific). Most sex-specific gene associations were identified in females (73%) and associated with tau tangles and longitudinal cognition (90%). Four X-linked genes, *MCF2*, *HDAC8*, *FTX*, and *SLC10A3*, demonstrated significant sex differences in their associations with AD endophenotypes (i.e., significant *sex x gene* interaction). Our results also uncovered sex-specific biological pathways, including a female-specific role of neuroinflammation and neuronal development, reinforcing the potential for sex-aware analyses to enhance precision medicine approaches in AD.

## Main Text

Women are disproportionately affected by Alzheimer’s disease (AD); two-thirds of all clinical cases of AD dementia in the United States are women.^1^ Women exhibit greater vulnerability to AD neuropathology, with each additional unit of pathology increasing the odds of progressing to clinical AD by approximately 22-fold in women relative to 3-fold in men.^2^ Females also experience more rapid cognitive decline in the presence of AD pathology^3^ and present with higher levels of AD pathology at autopsy, even among those deemed clinically normal prior to death.^2, 4^ However, the molecular factors that underlie these sex disparities remain poorly understood. Large-scale omics studies provide a unique opportunity to uncover sex-specific molecular mechanisms of AD.

Previous transcriptome-wide studies have shown significant differential gene expression between the sexes in postmortem AD brains,^5–8^ and sex-biased molecular pathways, including neuroinflammation and bioenergetic metabolism, have been identified.^7^ However, gaps in knowledge remain. Few studies have explored sex-specific gene expression changes in relation to the core AD neuropathologies, β-amyloid (Aβ) plaques and tau neurofibrillary tangles, or the clinical presentation of the disease outside of case/control analyses. In addition, studies have frequently excluded sex chromosomes due, in part, to technical considerations resulting from pseudo-autosomal regions or X inactivation, as examples.^9^ Given the etiologic complexity of AD, examination of the entire transcriptome driving AD endophenotypes offers an invaluable opportunity to better understand the underlying, and potentially sex-specific, biological pathways driving disease.

Thus, we performed the largest and most comprehensive study of sex-specific transcriptomic associations with AD endophenotypes (i.e., Aβ plaques, tau tangles, and longitudinal cognition) to date leveraging bulk RNA sequencing (RNAseq) data from over 750 sex-matched decedents (spanning 1490 samples from the dorsolateral prefrontal cortex, posterior cingulate, and caudate nucleus) enrolled in the Religious Orders Study/Memory and Aging Project (ROS/MAP).^10–12^

## Results

### Study Cohort Characteristics

This study leveraged brain tissue samples taken at autopsy from ROS/MAP participants. Given the large sample size difference between males (n=328) and females (n=640), the sexes were propensity score matched using age of death, post-mortem interval (PMI), education, latency to death (i.e., difference between age at death and age at last cognitive visit), race, and *APOE-*ε4 allele count as criteria (see **Methods**) for each brain region: the dorsolateral prefrontal cortex (DLPFC), posterior cingulate cortex (PCC), and head of the caudate nucleus (CN). The final analytical dataset included 1490 samples across 767 unique decedents, whose characteristics are presented by sex and brain region in **Table 1**. Overall, the mean age at death of the entire sample was 88.3 years (Standard Deviation (SD)=6.5, Range=67.3-108.3), and the average number of annual follow-up visits prior to death was 7.4 (SD=4.9, Range=0-23) (**Supplementary Table 1**). Most participants self-identified as non-Hispanic White (98%), and 26% of participants carried at least one copy of *APOE-*ε4. There were no statistically significant sex differences in criteria used for matching across any brain region (p>0.05). Notably, females had greater tau tangle burden in comparison to males in all brain regions, following previous reports (p<0.05, **Table 1**).^13–15^

**Table 1:** Participant Demographics by Brain Region.

|  | DLPFC |  |  | PCC |  |  | CN |  |  |
| --- | --- | --- | --- | --- | --- | --- | --- | --- | --- |
|  | Female | Male | P | Female | Male | P | Female | Male | P |
| **N** | 311 | 311 |  | 194 | 194 |  | 240 | 240 |  |
| **Age at Death (years)** | 87.63±6.55 | 87.51±6.61 | 0.82 | 88.19±6.51 | 87.84±6.31 | 0.59 | 87.42±6.35 | 87.53±6.43 | 0.85 |
| **Education (years)** | 16.90±3.24 | 17.30±3.85 | 0.17 | 16.76±3.03 | 17.14±3.96 | 0.29 | 16.80±3.25 | 17.35±3.80 | 0.09 |
| **Follow-up (years)** | 7.49±5.07 | 7.12±4.93 | 0.36 | 6.99±4.23 | 7.08±4.87 | 0.85 | 7.40±4.59 | 7.24±4.86 | 0.70 |
| **Latency to Death (years)** | 0.70±0.99 | 0.66±0.72 | 0.58 | 0.80±0.91 | 0.68±0.80 | 0.16 | 0.75±0.91 | 0.66±0.74 | 0.27 |
| **Post-mortem Interval (hours)** | 7.63±4.52 | 7.53±4.43 | 0.76 | 6.80±4.02 | 6.73±3.85 | 0.86 | 7.60±4.39 | 7.64±4.54 | 0.92 |
| **% Normal Cognition at Death** | 37 | 34 | 0.52 | 38 | 36 | 0.39 | 38 | 32 | 0.32 |
| **% APOE-ε4 Positive** | 25 | 25 | 0.90 | 27 | 27 | 0.99 | 26 | 26 | >0.99 |
| **% Non-Hispanic White** | 98 | 98 | 0.82 | 99 | 97 | 0.42 | 99 | 99 | 0.51 |
| **Amyloid (mm<sup>2</sup>)** | 3.90±4.20 | 3.69±3.97 | 0.55 | 3.81±4.09 | 3.94±4.07 | 0.77 | 4.18±4.06 | 3.94±4.18 | 0.52 |
| **Tau Tangles (mm<sup>2</sup>)** | **7.10±8.80** | **5.06±7.34** | **0.002** | **6.84±9.32** | **4.78±5.96** | **0.01** | **6.79±8.52** | **4.97±7.62** | **0.01** |
| **Global Cognition Score at Last Visit** | -0.69±1.10 | -0.72±1.01 | 0.75 | -0.62±1.05 | -0.67±0.99 | 0.64 | -0.61±1.03 | -0.71±1.04 | 0.34 |
| **Annual Change in Cognition** | -0.10±0.09 | -0.09±0.09 | 0.19 | -0.08±0.08 | -0.09±0.08 | 0.59 | -0.10±0.10 | -0.09±0.08 | 0.53 |

Values are presented as mean±standard deviation unless otherwise indicated.
Abbreviations: DLPFC=dorsolateral prefrontal cortex; PCC=posterior cingulate cortex, CN=caudate nucleus.
Cognition was operationalized as a global cognition composite, an average of z-scores from 17 tests across 5 domains of cognition (episodic, semantic, and working memory, perceptual orientation, and perceptual speed).
Amyloid and tau tangle density were measured via immunohistochemistry; values are quantified as the average area occupied by pathology across 8 brain regions (hippocampus, angular gyrus, and entorhinal, midfrontal, inferior temporal, calcarine, anterior cingulate, and superior frontal cortices).
P-values comparing male and females for each brain region were calculated using two-sided t-tests or chi-squared tests where applicable. Traits with p<0.05 are in bold.

### Sex-specific gene expression signatures of AD neuropathology and clinical presentation

To identify sex-specific gene expression signatures of AD, we conducted both sex-stratified (i.e., male and female analyses separately) and *sex x gene* interaction analyses. We examined brain gene expression associations with continuous immunohistochemical measures of Aβ and tau burden at autopsy and with a longitudinal global cognition composite score. Expression of 17,091 autosomal, 645 X-linked genes, and 83 Y-linked genes derived from RNAseq of bulk brain tissue were tested using linear regression models for cross-sectional autopsy outcomes and mixed effect models for the longitudinal cognition outcome. For sex-stratified analyses, all models included age at death and post-mortem interval as covariates, while longitudinal cognitive models additional included years of education and latency to death. Y-linked analyses were only conducted in males. False discovery rate (FDR) correction was performed for genes expressed on autosomal chromosomes, the X chromosome, and Y chromosome separately so that smaller effects on sex chromosomes may be observed. Sex-specific gene associations were defined as related to a trait in one sex (p_FDR_ ≤ 0.05) but not in the other sex in stratified analyses and showed suggestive evidence of a sex-modifying effect in interaction analyses (*sex x gene* interaction p<0.05, uncorrected). Significance for *sex x gene* interaction analyses for autosomal genes and X-linked genes was set *a priori* at p_FDR_ ≤ 0.10. Results from sensitivity and *post hoc* analyses (i.e. three-way interactions with *APOE*-ε4) were not corrected for multiple comparisons. FDR-corrected p-values will be notated as “p_FDR_” throughout; “p” will be used for uncorrected p-values.

Transcriptome-wide analyses yielded over 23,118 significant associations with Aβ, tau tangles, and longitudinal cognition (22,489 autosomal, 629 X-linked, and 0 Y-linked). Approximately 10% (2,320 of 23,118) of associations were sex-stratified, significant in one sex but not the other, as illustrated in **Figure 1**. In general, the pattern of sex-specific associations was consistent across the DLPFC, PCC, and CN and both the autosomal and X chromosomes (**Supplementary Figure 1**). In all three brain regions, we identified a total of 1697 female-stratified (1622 autosomal, 75 X-linked) and 623 male-stratified (609 autosomal, 14 X-linked) gene associations with Aβ, tangles, and longitudinal cognition (**Figure 1**). Most associations were identified in the DLPFC (70%, 1,619 of 2,320 associations) and with tau tangles (39%, 908 of 2,320) or longitudinal cognition (52%, 1182 of 2,320), **Supplementary Table 2**).

**Figure 1:**
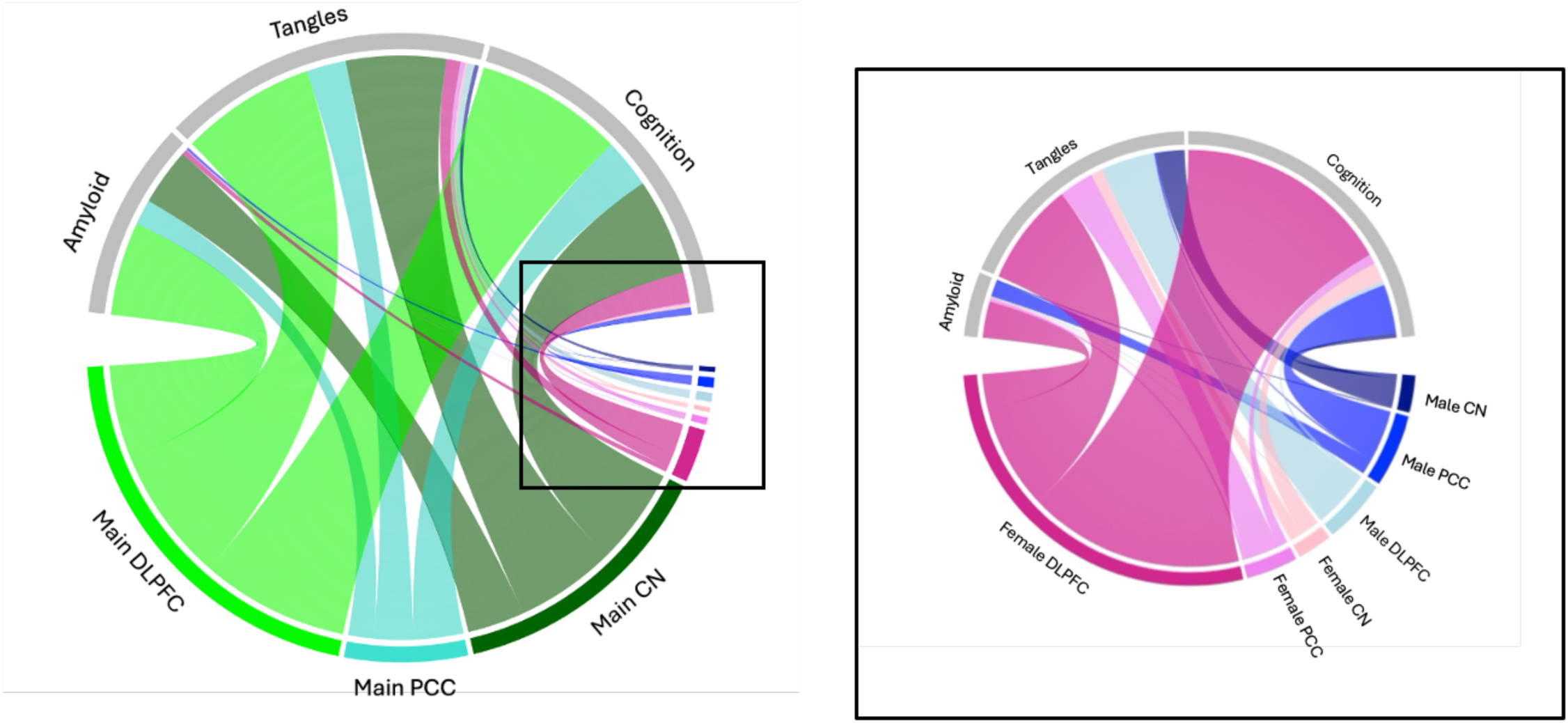
Sex-specific gene associations are roughly 10% of all significant gene associations identified. Chord diagrams summarizing the relative number of both autosomal and X-linked genes associated with Aβ, tau tangles, or longitudinal cognition in the Dorsolateral prefrontal cortex (DLPFC) or Posterior Cingulate Cortex (PCC) or Caudate Nucleus (CN) with chord size corresponding to number of significant genes. The right panel further highlights the significant male- and female-specific gene associations in sex-stratified analyses.

No genes showed significance across all three brain regions. **Table 2** highlights in bold those sex-stratified associations observed in at least two brain regions and demonstrated consistent directions of effect. Featured in this list are six male-stratified autosomal gene associations with Aβ (*GSR*) and tau tangles (*C16orf74*, *TDH-AS1*, *PNPLA6*, *ATG4B*, *Novel Transcript* (ENSG00000262691)); **Table 2**). Greater expression of *GSR*, *TDH-AS1*, and *C16orf74* were associated with lower levels of neuropathology, whereas *ATG4B*, ENSG00000262691, and *PNPLA6* were associated with higher pathological burden. *GSR*,^16^ *PNPLA6*,^17, 18^ and *ATG4B*^19^ have previously been implicated in neurodegenerative disorders, including AD. Additionally, two female-stratified autosomal genes were associated with tau tangles consistently across 2 brain regions (*MYO15A*, *NNT*), and a further 7 genes with longitudinal cognition (*PEPD*, *CTNNBL1*, *MVB12A*, *XKR8*, *WBP1L*, *GALNT7-DT*, *GCA*; **Table 2**). Higher levels of *NNT*, *MYO15A*, and *GCA* were protective in females; greater expression of *PEPD*, *CTNNBL1*, *MVB12A*, *XKR8*, *WBP1L*, and *GALNT7-DT* were associated with faster cognitive decline in females. *MYO15A*,^20^ *NNT*^21^ and *GCA*^22^ have previously been implicated in AD. *XKR8*^23, 24^ and *WBP1*L^25^ are related to other disorders, such as schizophrenia, and *CTNNBL1* has previously been associated with memory in a young, cognitively healthy cohort.^26^

**Table 2:** Sex-Specific Autosomal Associations with Consistent Effects Across Two of Three Brain Regions.

| Male-specific Gene Associations |  |  |  |  |  |  |  |  |  |  |  |  |  |  |
| --- | --- | --- | --- | --- | --- | --- | --- | --- | --- | --- | --- | --- | --- | --- |
|  |  |  | DLPFC |  |  |  | PCC |  |  |  | CN |  |  |  |
| Outcome | Gene | Chr | B | SE | T | P<sub>FDR</sub> | B | SE | T | P<sub>FDR</sub> | B | SE | T | P<sub>FDR</sub> |
| Amyloid | GSR | 8 | -0.04 | 0.06 | -0.68 | 0.83 | -0.29 | 0.08 | -3.68 | **0.04** | -0.34 | 0.07 | -4.77 | **0.01** |
| Tau Tangles | ATG4B | 2 | 0.16 | 0.05 | 3.31 | **0.03** | 0.03 | 0.06 | 0.48 | 0.89 | 0.16 | 0.05 | 3.05 | **0.05** |
|  | TDH-AS1 | 8 | -0.22 | 0.05 | -4.42 | **0.002** | -0.14 | 0.06 | -2.48 | 0.24 | -0.23 | 0.06 | -3.98 | **0.01** |
|  | C16orf74 | 16 | -0.16 | 0.05 | -3.29 | **0.03** | 0.08 | 0.06 | 1.22 | 0.63 | -0.25 | 0.06 | -4.27 | **0.004** |
|  | ENSG00000262691 | 16 | 0.15 | 0.05 | 2.97 | **0.05** | 0.05 | 0.06 | 0.82 | 0.77 | 0.17 | 0.05 | 3.18 | **0.04** |
|  | PNPLA6 | 19 | 0.17 | 0.05 | 3.43 | **0.02** | 0.11 | 0.06 | 1.90 | 0.40 | 0.22 | 0.06 | 3.81 | **0.01** |
| Female-specific Gene Associations |  |  |  |  |  |  |  |  |  |  |  |  |  |  |
|  |  |  | DLPFC |  |  |  | PCC |  |  |  | CN |  |  |  |
| Outcome | Gene | Chr | B | SE | T | P<sub>FDR</sub> | B | SE | T | P<sub>FDR</sub> | B | SE | T | P<sub>FDR</sub> |
| Tau Tangles | NNT | 5 | -0.21 | 0.05 | -3.99 | **0.01** | -0.31 | 0.07 | -4.74 | **0.005** | -0.05 | 0.06 | 0.80 | 0.72 |
|  | MYO15A | 17 | -0.21 | 0.05 | -3.95 | **0.01** | -0.07 | 0.07 | -0.96 | 0.70 | -0.22 | 0.06 | -3.69 | **0.03** |
| Longitudinal Cognition | XKR8 | 1 | -0.03 | 0.01 | -4.21 | **0.001** | -0.01 | 0.01 | -1.03 | 0.70 | -0.04 | 0.01 | -4.07 | **0.01** |
|  | GCA | 2 | 0.01 | 0.01 | 1.92 | 0.16 | 0.03 | 0.01 | 3.76 | **0.04** | 0.03 | 0.01 | 3.10 | **0.05** |
|  | GALNT7-DT | 4 | -0.03 | 0.01 | -4.08 | **0.001** | 0.01 | 0.01 | 0.56 | 0.86 | -0.03 | 0.01 | -3.24 | **0.04** |
|  | WBP1L | 10 | -0.02 | 0.01 | -3.73 | **0.004** | -0.01 | 0.01 | -0.56 | 0.86 | -0.03 | 0.01 | -3.27 | **0.04** |
|  | MVB12A | 19 | -0.03 | 0.01 | -3.94 | **0.002** | -0.02 | 0.01 | -2.73 | 0.17 | -0.04 | 0.01 | -4.40 | **0.003** |
|  | PEPD | 19 | -0.03 | 0.01 | -4.10 | **0.001** | -0.01 | 0.01 | -1.42 | 0.57 | -0.03 | 0.01 | -4.65 | **0.002** |
|  | CTNNBL1 | 20 | -0.02 | 0.01 | -3.08 | **0.02** | -0.03 | 0.01 | -4.04 | **0.02** | -0.02 | 0.01 | -2.01 | 0.25 |

Significant values are bolded.
Number of genes in each brain region: DLPFC (n=926, genes=17,088 (autosomal), 562 (X-linked), 63 (Y-linked)); PCC (n= 518, genes=18,385 (autosomal), 598 (X-linked), 71 (Y-linked)); CN (n=706, genes=17,928 (autosomal), 591 (X-linked), 76 (Y-linked)).
Abbreviations: DLPFC=dorsolateral prefrontal cortex; PCC=posterior cingulate cortex, CN=caudate nucleus.

To determine whether *APOE*-ε4, the strongest known common genetic risk factor for AD, played a role in our sex-stratified associations with Aβ or tau tangles, we performed sensitivity analyses additionally covarying for *APOE*-ε4 allele status. All sex-stratified gene associations with Aβ and tau in two brain regions remained significant. Similarly, models in which cognition was the outcome were additionally covaried for Aβ or tau burden, as pathologic burden is known to independently effect cognition. All sex-stratified gene associations with cognition in two brain regions remained significant in sensitivity analyses (**Supplementary Table 3**).

### Sex modifies transcriptome associations with AD endophenotypes

We examined *sex x gene* interaction models to directly investigate sex differences in gene associations. At a threshold of p_FDR_ ≤ 0.10, sex significantly moderated four X-linked genes (*MCF2*, *HDAC8*, *SLC10A3*, and *FTX*) on tau tangle burden or cognitive decline in the DLPFC (**Table 3**, **Figure 2**). Higher expression of *MCF2* was significantly associated with lower levels of tau tangles in females (β=-0.18, p_FDR_=0.02, **Figure 2A**), but not in males (β=0.10, p_FDR_=0.26). By contrast, higher expression of *HDAC8* (**Figure 2B**) was associated with higher tau tangle burden in females (β=0.22, p_FDR_=0.01), but not in males (β=-0.09, p_FDR_=0.37). Higher *SLC10A3* (**Figure 2C**) expression was associated with faster cognitive decline in females (β=-0.02, p_FDR_=0.04), but not in males (β=0.02, p_FDR_=0.24). Alternatively, *FTX* (**Figure 2D**) expression was significantly associated with slower cognitive decline in males (β=0.02, p_FDR_=0.05), while no relationship was found in females (β=-0.02, p_FDR_=0.13). No other *sex x gene* interaction, whether autosomal or X- or Y-linked, survived FDR-correction for multiple comparisons.

**Figure 2:**
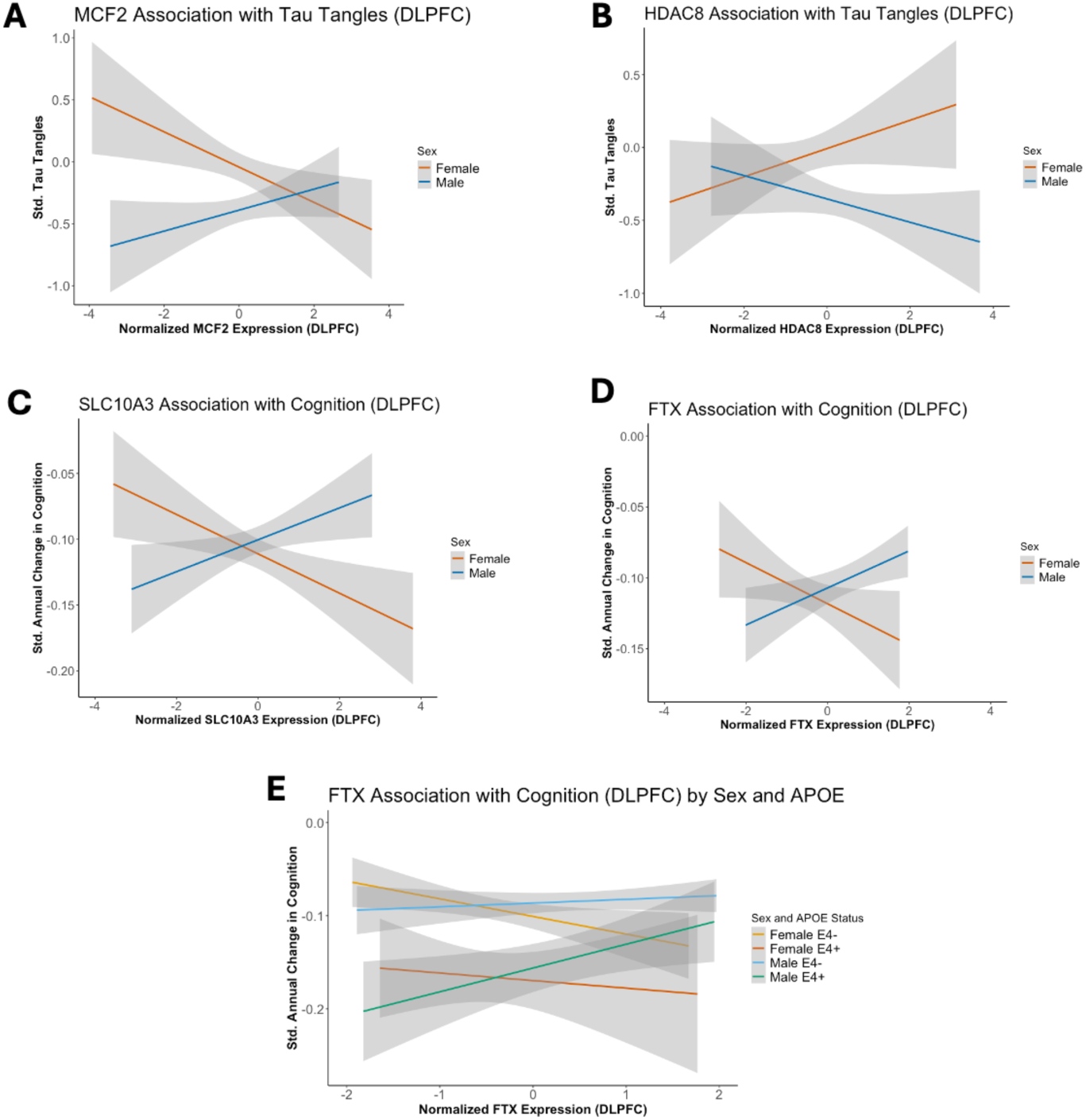
*MCF2*, *HDAC8*, *SLC10A3*, and *FTX* associations with tau tangles and cognition are moderated by sex. A-D) Plots demonstrating the relationship between gene expression (x-axis) and AD endophenotype (y-axis), and shaded regions represent the 95% confidence interval. Females are colored in orange and males are colored in blue. E) A plot demonstrating the relationship between *FTX* and cognitive decline colored by *APOE*-ε4 status and sex. Females are colored in yellow (ε4-) and orange (ε4+); males are colored in green (ε4-) and blue (ε4+). Shaded regions represent the 95% confidence interval.

**Table 3:** Significant Sex x X-Linked Gene Interactions in the DLPFC.

| Outcome | Gene | Analysis | $\beta$ | SE | T | P<sub>FDR</sub> |
| --- | --- | --- | --- | --- | --- | --- |
| Tau Tangles | *MCF2* | **Sex x *MCF2*** | **0.28** | **0.07** | **3.74** | **0.052** |
|  |  | Male-stratified | 0.10 | 0.05 | 1.94 | 0.26 |
|  |  | **Female-stratified** | **-0.18** | **0.06** | **-3.33** | **0.02** |
|  | *HDAC8* | **Sex x *HDAC8*** | **-0.30** | **0.08** | **-3.82** | **0.052** |
|  |  | Male-stratified | -0.08 | 0.05 | -1.60 | 0.37 |
|  |  | **Female-stratified** | **0.22** | **0.06** | **3.75** | **0.01** |
| Longitudinal Cognition | *SLC10A3* | **Sex x *SLC10A3*** | **0.04** | **0.01** | **3.69** | **0.06** |
|  |  | Male-stratified | 0.02 | 0.01 | 2.51 | 0.24 |
|  |  | **Female-stratified** | **-0.02** | **0.01** | **-2.7.** | **0.04** |
|  | *FTX* | **Sex x *FTX*** | **0.04** | **0.01** | **3.82** | **0.06** |
|  |  | **Male-stratified** | **0.02** | **0.01** | **3.58** | **0.0496** |
|  |  | Female-stratified | -0.02 | 0.01 | -2.06 | 0.13 |

Significant values are bolded.

Similar to the sex-stratified models, the interaction models for *MCF2*, *HDAC8*, *SLC10A3*, and *FTX* were additionally adjusted for *APOE-ε4* allele positivity and/or AD neuropathology, where relevant, to see whether the genes had effects on tau tangles and cognition independent of *APOE-ε4* and neuropathology. All four associations remained significant when covarying for *APOE-ε4* status and neuropathological burden (**Supplementary Table 4**) suggesting that these genes do have an additional impact on AD endophenotypes beyond *APOE-ε4* and neuropathology. The sex-stratified associations for these four genes also remained significant in sensitivity analyses.

### Differential expression of gene x sex interaction candidates

One challenge of studying the X chromosome is the accounting for X chromosome inactivation, a biological process during which one X chromosome in females is randomly silenced to prevent gene overdosing in females in comparison to males. Due to this phenomenon, we wanted to examine whether the moderating effects of sex on *MCF2*, *HDAC8*, *SLC10A*, and *FTX* were due to sex differences in gene expression between males and females. *FTX* and *HDAC8* were expressed more highly in males than females (Wilcoxon p<0.0001), *SLC10A3* was expressed more highly in females than males (Wilcoxon p=0.05), and *MCF2* expression did not significantly differ between sexes (**Supplementary Figure 2**).

### APOE-ε4 further modifies the interaction between sex and gene expression on tau tangles and cognition

Sex differences in the effect of *APOE*-ε4 on AD risk and on AD endophenotypes are well-established.^3, 14^ Therefore, we performed *post hoc* analyses examining the possible modifying effects of both sex and *APOE*-ε4 for the four X-linked genes (*MCF2*, *HDAC8*, *SLC10A*, and *FTX*) by leveraging a three-way interaction model. We observed a significant *APOE*-ε4 *x sex x FTX* interaction on longitudinal cognition (p=0.03, **Figure 2E**, **Supplementary Table 5**), whereby the male-specific protective effect of *FTX* expression on cognitive decline was driven by *APOE*-ε4 carriers (β=0.05, p=0.01, **Figure 2E**).

### Exploratory analysis of the Y Chromosome

In males, we also examined the association between 83 Y-linked genes (**Supplementary Table 6**) and Aβ, tau tangles, and longitudinal cognition in the DLPFC, PCC, and CN. No genes survived correction for multiple comparisons at a p_FDR_ ≤ 0.05.

### Post-hoc pathway analysis of sex-specific associations

To connect our sex-stratified findings to broader biological mechanisms that may be implicated in the neuropathological progression of AD in one sex but not the other, we completed pre-ranked gene set enrichment analyses. Pathway enrichment was performed for 18 autosomal gene lists; one list per sex, outcome, and brain region (e.g., female-specific, DLPFC, Aβ). All tested autosomal genes were included in these gene lists (DLPFC = 17,088, PCC = 18,385, CN = 17,928). Gene Ontology: Biological Process (GO:BP) terms between the size of 15 to 500 genes were used for annotation; genes were ranked based on the beta coefficient from sex-stratified linear regression models for each outcome and brain region. Results were FDR-corrected to account for multiple comparisons (p_FDR_ ≤ 0.05). Top associated pathways are presented in **Figure 3**; all gene set enrichment results are presented in **Supplementary Table 7**.

**Figure 3:**
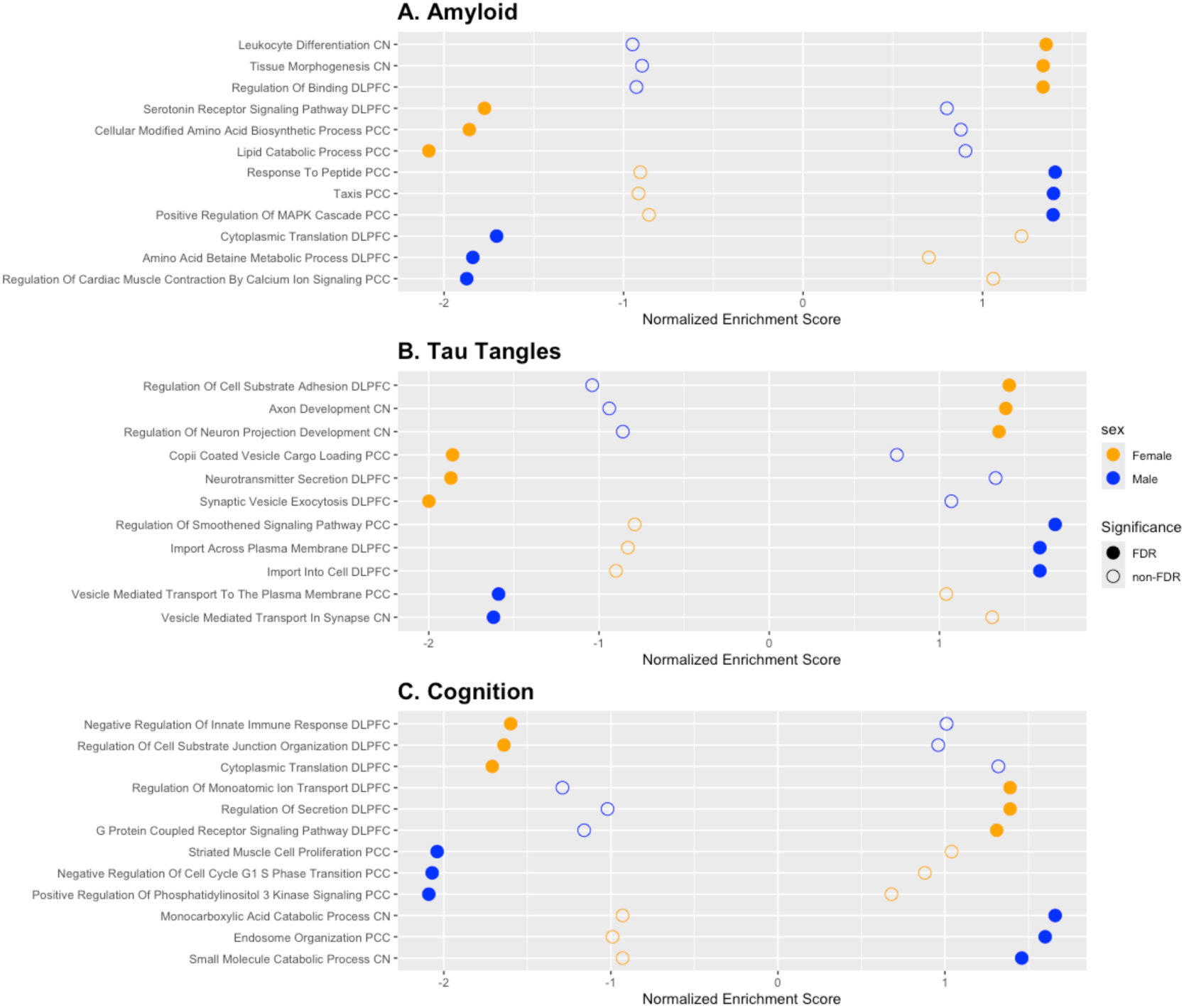
Significant GO Biological Progress terms differ between males and females. Dotplots demonstrating GO:BP terms for each AD endophenotype. Depicted terms were selected using the following criteria: An FDR-corrected p-value significant in one sex, but not the other, and opposite signs of normalized enrichment scores (NES). The plots include the top 3 highest and lowest enrichment scores for each sex. Females are denoted by yellow points, and males in blue. Filled circles represent a significant FDR-corrected p-value while an empty circle does not. Brain regions are included at the end of each GO:BP term: Dorsolateral prefrontal cortex (DLPFC) or Posterior Cingulate Cortex (PCC) or Caudate Nucleus (CN).

Pathway enrichment analyses yielded a total of 4,190 significantly enriched GO:BP pathways (1,589 unique) terms across all examined gene lists; most enriched pathways arose from the DLPFC (2,234 of 4,190). Female-stratified analyses resulted in 2,774 enriched pathways, in contrast with 1,416 for males. Pathways related to amyloid and tau tangles were most enriched in females (2,193 of 2,774) in comparison with cognition, whereas more GO:BP pathways related to cognition were enriched in males (767 of 1,416). 676 of 4,190 significant terms overlapped between sexes; 390 of those terms were related to males and females in opposite directions (i.e., upregulated for one sex, downregulated for the other).

When more closely examining sex-specific enriched pathways, genes involved in neuronal development and function, along with innate immunity, were upregulated in females in response to AD neuropathology whereas neurotransmission and cellular metabolism-related pathways were downregulated in females (**Figure 3**). Cellular growth and migration, as well as endocytosis-related pathways, were primarily upregulated in males in response to amyloid and tau pathology.

## Discussion

Approximately 10% of our observed autosomal and X-linked gene expression associations with AD endophenotypes exhibited sex-stratified significance. 89 of the 2,320 (3.8%) sex-specific associations were X-linked, which is consistent with the proportion of protein-coding genes (4% X-linked) within the entire genome.^27^ Most sex-specific associations were female-stratified (73.1%) and were predominantly associated with post-mortem tau tangles and ante-mortem cognitive trajectories. The direction of effects suggested increased gene expression was associated with both risk for and protection against AD. Additionally, our study is one of the first to demonstrate the lack of associations between Y-linked genes and AD endophenotypes.

Most of the sex-specific associations, whether involving stratified or interaction models, were with tau and cognition. Women at all stages of disease have greater tau burden than men as measured at autopsy, via tau-PET and cerebrospinal fluid (CSF) measures,^13, 14^ and this difference in tau burden is even more apparent if they carry *APOE*-ε4^3, 13, 28^ or have abnormal amyloid burden.^13^ Studies also demonstrate that women experience faster cognitive decline in response to tau pathology, especially if they are Aβ-positive and carry *APOE*-ε4.^3, 29^ Taken together, these findings provide evidence for transcriptomic pathways that underlie the well-characterized clinical findings of sex differences in AD risk.

Four X-linked genes, *MCF2*, *HDAC8*, *SLC10A3*, and *FTX*, displayed evidence of moderation by sex. These genes conferred both risk and protection, with greater *MCF2* and *FTX* expression associated with less tau tangles in females and slower cognitive decline in males, respectively. Further, protective *FTX* effects appeared to be driven by male *APOE*-ε4 carriers. By contrast, greater *HDAC8* and *SLC10A3* expression was associated with greater tau tangle burden and faster cognitive decline, respectively, in females only. These four X-linked associations survived additional adjustment from *APOE*-ε4, the most common genetic risk factor for sporadic AD dementia suggesting these genes have an effect beyond *APOE*-ε4 carriership.

There has been no previous association between *MCF2* and Alzheimer’s disease endophenotypes, though it has been associated with autism spectrum disorders^30^ and cortical neuron migration.^31^ *HDAC8* and *SLC10A3* are members of protein families that have been implicated in AD.^32, 33^ *HDAC8* encodes a histone deacetylase; epigenetic deacetylation has been associated with AD.^32, 34^ Similarly, *SLC10A3* is a sodium-bile acid cotransporter and thought to be a housekeeping gene due to its ubiquitous expression;^35^ higher levels of bile acids have been observed in individuals with AD and mild cognitive impairment (MCI).^36^ Recently, another member of the SLC super family, *SLC9A7* was identified by Belloy et al., as a risk locus in an X-chromosome wide association study.^37^ Though we see a protective effect in males, *FTX* is a long non-coding RNA that is heavily involved in X chromosome inactivation,^38^ which males do not undergo, warranting further study. We posit that some of the protective effects seen on X-linked genes for men, particularly on *FTX*, may be due to a demographically skewed sample. However, these protective effects could also be observed on the maternally inherited X chromosome.^39^ In addition, both *FTX* and *HDAC8* showed higher expression in males compared to females despite being X-linked and related to X inactivation,^38, 40^ though their expression may not be increased compared to females in other datasets and samples requiring further investigation. These findings support a synergistic role for the X chromosome in AD and further highlights the importance of including it in future studies.

Though some pathways overlap between the two sexes, many of the biological pathways enriched in our study differed between males and females underscoring the thesis of distinct sex-specific pathways that may lead each sex to higher risk of AD. Females exhibited upregulation in neuronal development and immune-related pathways. A study by Meyer et al. also identified an upregulation of genes involved in neurogenesis in induced pluripotent stem cells derived from individuals with sporadic AD, supporting our findings.^41^ They suggested that premature neuronal differentiation as a result of this overexpression may contribute to the onset of AD. Additionally, neuroinflammation and innate immunity has been implicated in the development of AD.^42–44^ Females also exhibited downregulation in neurotransmission-related pathways. Disruption in neurotransmission has long been linked to AD, perhaps resulting from neuronal death during the progression of disease.^45^ In contrast, males exhibited upregulation in endocytosis pathways. Cellular aging is suggested to facilitate increased amyloid precursor protein (APP) uptake, resulting in the creation of more Aβ deposits.^46, 47^ Similarly, tau hyperphosphorylation has been linked to aberrant endocytosis, both of which may contribute to the development of AD.^46^ Our study also highlighted changes to the cell cycle in males, described in the two-hit hypothesis of AD^48^ such that abnormal neuronal reentry into the cell cycle results in cell death.^49^ We further highlight biological processes that are upregulated in one sex, and downregulated in the other (**Figure 3**) perhaps suggesting that one intervention may not be effective in both sexes. These data may point toward the development of precision therapeutics that may function differently for each sex on the same pathway.

Our study is the largest transcriptomic investigation of the X chromosome and AD endophenotypes to date. We identified 4 gene associations that were moderated by sex, and 89 other X-linked gene associations were sex-stratified. One main technical challenge is that X chromosome dosage may influence the expression of X-linked genes. In females, one X chromosome is silenced at random to equalize dosage between males and females.^50^ The silencing is not complete; up to a third of genes escape inactivation, which can result in increased expression in females.^51, 52^ 40 of our significant female-specific genes have been previously reported to be X-inactivation escapees, such as *TSPAN6*^53^ and *SEPTIN6*,^54, 55^ which have also been implicated in AD (**Supplementary Table 8**). The upregulation of immune-related pathways in females warrants further study as many inflammatory genes are X-linked,^56^ and genes escaping X-inactivation have been linked to autoimmune disorders in females.^57^ This study had many strengths, including a large sample size, the examination of multiple brain regions, and gene expression from both X- and Y-chromosomes.

Nevertheless, this study also possessed some weaknesses. Most individuals in the sample identify as non-Hispanic white and are highly educated, which may not be generalizable to a larger population. In addition, participants were, on average, 88 years at their time of death. At such an advanced age, it is very possible that brain-derived gene expression might be indicating different disease processes than what might be observed *in vivo* at younger ages. With these types of data, we are also limited by the correlational nature of the neuropathological analyses. That is, one cannot infer causality or timing of effects. Further, we age-matched females and males in this sample, which may result in a male sample who is less representative of the general population, given the average male life expectancy is 73.5 years as of 2021.^58^ Given the increase in AD risk attributed to *APOE*-ε4 carriership,^59^ male *APOE*-ε4 carriers living to an average age of 88 likely required a considerable level of gene-environment protective effects to survive into advanced age.^60^ Taken together, similar studies in the future should consider a sample that is more representative of the general population.

Our study is the largest brain transcriptomic study of sex differences in AD and provides a comprehensive look at the autosome and sex chromosomes. We identified 2,320 sex-specific transcriptomic associations with amyloid, tau, and longitudinal cognition, with 89 associations on the X chromosome, and four of whom are significant moderators. Our findings support the involvement of processes such as endocytosis, neuronal development, and neurotransmission in the progression and development of AD, and suggest that biological pathways to AD may differ by sex.

Though we have successfully identified sex-specific gene associations with AD endophenotypes, we have yet to pinpoint cell-type specific mechanisms and assess the impact of X-chromosome inactivation escapism on our observed associations. Since these contributions remain unclear, the present study forms a strong foundation for future work with longitudinal and cell-specific data, which are critical to continue characterizing and uncovering sex differences in AD. Overall, these findings highlight the importance of precision medicine approaches that consider sex-specific biological pathways to select new targets for AD therapeutic intervention.

## Online Methods

### Study Population

Data were obtained from the Religious Orders Study and Rush Memory and Aging Project (ROS/MAP). These studies enrolled older adults free of dementia who agreed to annual clinical evaluations and brain donation at death.^10^ All participants gave written informed consent. The Rush Institutional Review Board (IRB) approved all protocols and the Vanderbilt University Medical Center IRB approved secondary analyses of existing data. Data from this cohort is accessible either online on the Accelerating Medicines Partnership – Alzheimer’s Disease (AMP-AD) Knowledge Portal (https://adknowledgeportal.synapse.org/Explore/Programs/DetailsPage?Program=AMP-AD, syn3219045) or via the Rush Alzheimer’s Disease Center Resource Sharing Hub (https://www.radc.rush.edu/).

### Neuropsychological Composite Scores

Cognitive measures in ROS/MAP have been described previously.^61^ A global cognition composite was calculated by averaging z-scores from 17 tests across 5 domains of cognition (episodic, semantic, and working memory, perceptual orientation, and perceptual speed).

### Bulk mRNA Sequencing

Dorsolateral prefrontal cortex (DLPFC), posterior cingulate cortex (PCC), and the head of the caudate nucleus (CN) tissue were acquired at autopsy. These regions were selected by the parent study based on tissue availability and because the regions are affected by numerous age-related disease conditions. Full details on RNA extraction, library preparation, and RNA sequencing have been described previously.^11, 12^ Briefly, RNA was extracted from each brain region and ribosomal depletion was formed (RiboGold (Illumina, 20020599). Sequence libraries were sequenced using 2 x 150bp paired end reads on an Illumina NovaSeq 6000 targeting an average of 40 to 50 million reads.

### RNAseq Alignment and Quality Control

Sequence alignment and gene counting followed a published protocol^62, 63^ including STAR (version 2.5.2b)^64^ alignment to the Ensembl (GRCh38 release 99)^65^ reference genome. Gene counts were obtained using the featureCounts function from the Subread package (v.2.0.0).^66^ Alignment metrics were calculated using Picard tools (version 2.18.27, http://broadinstitute.github.io/picard/).^67^ Prior to quality control, the number of samples was as follows: n_DLPFC_ = 1,126, n_PCC_ = 531, n_CN_ = 726.

For quality control (QC) after alignment, samples with RNA integrity number (RIN) <4 or post-mortem interval (PMI) >24 hours were removed. At this time, genes were split into three groups by chromosome: autosomal (chromosomes 1-22), X, and Y chromosome (males only) and underwent QC separately. Prior to quantile normalization using the cqn package (version 1.48.0),^68^ genes with <1 count per million (CPM) in 50% of individuals per diagnosis (i.e., normal cognition, mild cognitive impairment (MCI), AD) were removed. Additionally, genes missing gene length or GC-content measurements were removed. After quantile normalization, gene expression values with >5 standard deviations (SD) from the mean were set to 0. Samples that were principal component outliers (greater than 5SD from the mean), whose principal components did not align to participants’ self-reported sex, or who were missing RIN or demographic variables (e.g., age, sex, cognition), were also removed. To confirm observed effects were not due to technical variation, RNAseq data underwent further iterative normalization using the *limma* package (version 3.60.4)^69, 70^ adjusting for batch, sex, age at death, post-mortem interval (PMI), RIN, and percentage of coding, intronic, and intergenic bases resulting in the following datasets: DLPFC [n=926, genes=17,088 (autosomal), 562 (X-linked), 63 (Y-linked)]; PCC [n= 518, genes=18,385 (autosomal), 598 (X-linked), 71 (Y-linked)]; CN [n=706, genes=17,928 (autosomal), 591 (X-linked), 76 (Y-linked)]. A visual workflow including the number of samples dropped at each step is included as **Supplementary Figures 3-4**.

### Measures of Alzheimer’s Disease Pathology

Aβ load and tau tangle density were measured via immunohistochemistry, as detailed previously.^2^ Aβ load (cortical) and tau tangle density was quantified as the average area occupied by pathology across 8 brain regions: hippocampus, angular gyrus, and entorhinal, midfrontal, inferior temporal, calcarine, anterior cingulate, and superior frontal cortices. Aβ and tau tangle values were square-root transformed to approximate a normal distribution for analyses.

### Statistical Analyses

Statistical analyses were completed using R (version 4.2.1). Due to a larger number of female participants, we subsampled female participants by matching to male participants using propensity scores based on age of death, PMI, education, latency to death (i.e., difference between age of death and age at last cognitive visit), race, and *APOEε4* allele count. After matching, males and females were statistically compared across these measures to ensure no differences remained between groups. Gene expression, Aβ, tau tangles, and cognitive measures were mean-centered prior to analyses and were treated as continuous variables. Linear regression and mixed-effects models were used to assess the association between gene expression and AD endophenotypes for each gene in both pooled and sex-stratified analyses. All p-values for discovery analyses were adjusted using the false discovery rate (FDR) method to correct for multiple comparisons across each model, brain region, sex-stratification, and outcome. FDR-correction was performed for genes expressed on autosomal chromosomes, the X chromosome, and Y chromosome separately so that smaller effects on sex chromosomes may be observed. Significance in sex agnostic (i.e., pooled) analyses was set *a priori* to FDR-corrected p ≤ 0.05. Sex-specific gene associations were defined as related to a trait in one sex (p_FDR_ ≤ 0.05) but not in the other sex in stratified models and showed suggestive evidence of a sex-modifying effect in interaction models (*sex x gene* interaction p<0.05, uncorrected). *Gene x sex* interaction significance was set *a priori* to p_FDR_ ≤ 0.10. Age at death, PMI and sex were included as covariates where relevant (i.e., sex in male-female pooled analyses) in the cross-sectional models examining Aβ load and tau tangle density. Linear mixed-effects models tested the association between gene expression and longitudinal global cognition where the intercept and time interval between the final cognitive visit and the visit of interest were entered as both fixed and random effects. Mixed-effects models included age at death, PMI, latency to death, education, and sex (where relevant) as covariates.

*APOE-*ε4 status (positive/negative) was included as an additional covariate in sensitivity analyses; neuropathological measures were also added as covariates when cognition was used as an outcome. A linear mixed-effects model including the aforementioned covariates was used to assess the three-way *sex x gene x APOE-ε4* status on cognition as a *post hoc* analysis. Sensitivity and *post hoc* analyses were not corrected for multiple comparisons. Wilcoxon tests were used to compare mean normalized gene expression of between males and females. FDR-corrected p-values will be notated as “p_FDR_” throughout; “p” will be used for uncorrected p-values.

### Gene Set Enrichment Analysis

We performed pre-ranked gene set enrichment analyses with the R package fgsea (version 1.26.0)^71^ on all tested autosomal genes per sex, brain region, and outcome. Analyses were limited to Gene Ontology: Biological Process (C5)^72, 73^ terms of 15 to 500 genes. Gene rankings were based on the sex-stratified beta coefficients from each linear regression model. All enrichment results were FDR-corrected to account for multiple comparisons.

## Supporting information

Supplemental Tables (Large)

## Acknowledgements

The results published here are in whole or in part based on data obtained from the AD Knowledge Portal (https://adknowledgeportal.synapse.org). Study data were provided by the Rush Alzheimer’s Disease Center, Rush University Medical Center, Chicago. Data collection was supported through funding by NIA grants P30AG10161 (ROS), R01AG15819 (ROSMAP; genomics and RNAseq), R01AG17917 (MAP), R01AG30146, R01AG36042 (5hC methylation, ATACseq), RC2AG036547 (H3K9Ac), R01AG36836 (RNAseq), R01AG48015 (monocyte RNAseq) RF1AG57473 (single nucleus RNAseq), U01AG32984 (genomic and whole exome sequencing), U01AG46152 (ROSMAP AMP-AD, targeted proteomics), U01AG46161 (TMT proteomics), U01AG61356 (whole genome sequencing, targeted proteomics, ROSMAP AMP-AD), the Illinois Department of Public Health (ROS/MAP), and the Translational Genomics Research Institute (genomic). Additional phenotypic data can be requested at www.radc.rush.edu.

Additional support includes DP2-AG082342, R00-AG061238, K01-AG049164, R01-AG059716, R01-AG061518, R21-AG05994, K12-HD043483, K24-AG046373, HHSN311201600276P, S10-OD023680, R01-AG034962, R01-NS100980, R01-AG056534, P30-AG010161, R01-AG15819, R01-AG17917, U01-AG46152, Vanderbilt Clinical Translational Science Award (UL1-TR000445), the Vanderbilt Memory and Alzheimer’s Center, and the Vanderbilt Alzheimer’s Disease Research Center (P20-AGAG068082).

## Ethics Declarations

The authors declare no competing interests.

