## Supplemental Materials (Methods and Tables) for "Sex-specific Associations of Gene Expression with Alzheimer’s Disease Neuropathology and Ante-mortem Cognitive Performance"

**Supplementary Information**

**Supplementary Figure 1.** Chord diagram summarizing X-transcriptome wide analyses

**Supplementary Figure 2.** Expression of *MCF2*, *HDAC8*, *FTX*, and *SLC10A3* in males and females

**Supplementary Figure 3.** Autosomal and X chromosome RNA sequencing quality control (QC) flow chart

**Supplementary Figure 4.** Y chromosome RNAseq QC flow chart

**Supplementary Table 1.** Pooled demographics for 767 ROS/MAP participants

**Supplementary Table 2.** Results of sex-stratified autosomal and X-linked analyses

**Supplementary Table 3.** Results of sex-stratified autosomal and X-linked analyses adjusting for *APOE*ε4-positivity or AD neuropathological burden

**Supplementary Table 4.** Results including sensitivity analyses adjusting for *APOE*ε4-positivity or AD neuropathological burden for *MCF2*, *HDAC8*, *FTX*, and *SLC10A3* gene interactions in the DLPFC

**Supplementary Table 5.** Results of the three-way *sex x X-linked gene x APOE-ε4* interaction analyses

**Supplementary Table 6.** Results of Y-linked gene associations with AD endophenotypes

**Supplementary Table 7.** Results of Gene Ontology: Biological Process gene set enrichment analyses for autosomal genes

**Supplementary Table 8.** List of significant (sex-stratified) X-linked genes reported to escape X inactivation

**Supplementary Methods:**

Summary of models used in regression analyses

**Supplementary Methods**

**Summary of models used in analyses:**

Model A1: Aβ OR tau tangles ~ Participant Age of Death + Sex + Post-mortem Interval + Gene

Model A2: Aβ OR tau tangles ~ Participant Age of Death + Post-mortem Interval + Gene*Sex

Model A3: Aβ OR tau tangles ~ Participant Age of Death + Post-mortem Interval + Gene

- Male- or female-stratified

Model A4: Aβ OR tau tangles ~ Participant Age of Death + Post-mortem Interval + Gene + *APOE*ε4 status

Model B1: Longitudinal Cognition ~ Participant Age of Death + Sex + Post-mortem Interval + Latency to Death + Education + Gene

Model B2: Longitudinal Cognition ~ Participant Age of Death + Post-mortem Interval + Latency to Death + Education + Gene*Sex

Model B3: Longitudinal Cognition ~ Participant Age of Death + Post-mortem Interval + Latency to Death + Education + Gene

- Male- or female-stratified

Model B4: Longitudinal Cognition ~ Participant Age of Death + Post-mortem Interval + Latency to Death + Education + Gene + Amyloid + Tau Tangles

**Supplemental Figure 1:** Chord diagram summarizing X-transcriptome wide analyses

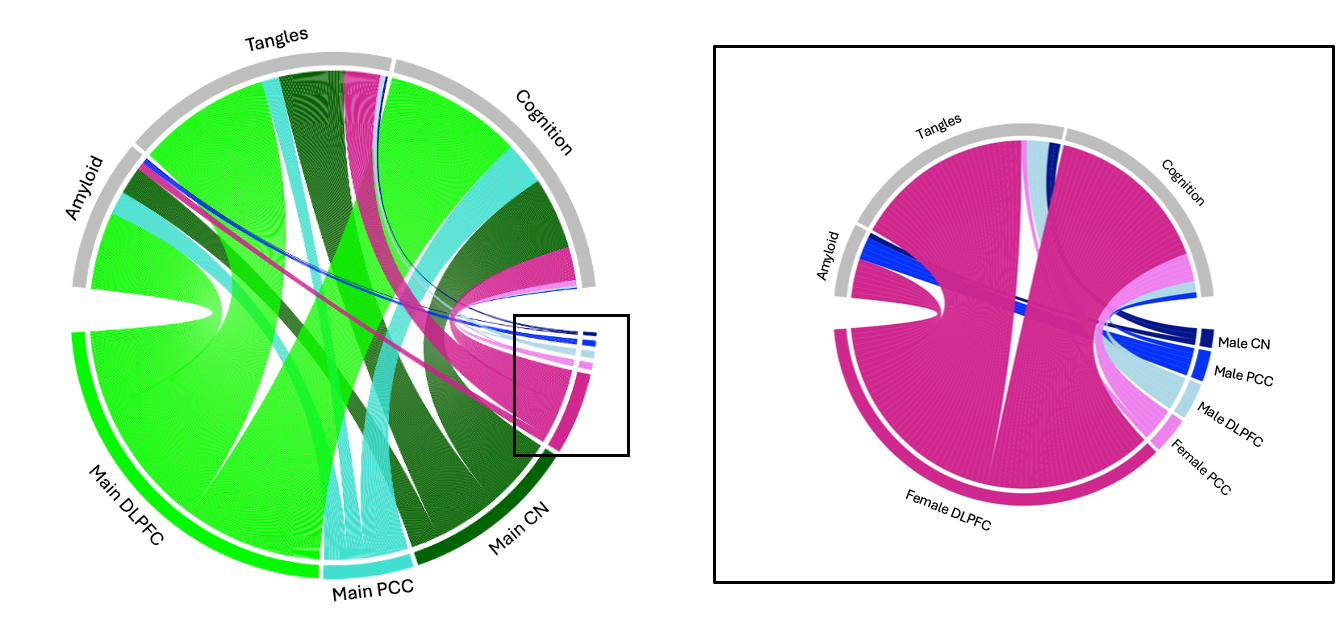

Chord diagrams summarizing the relative number of X-linked genes associated with Aβ, tau tangles, or longitudinal cognition in the Dorsolateral prefrontal cortex (DLPFC) or Posterior Cingulate Cortex (PCC) or Caudate Nucleus (CN) with chord size corresponding to number of significant genes. The right panel further highlights the significant male- and female-specific gene associations in sex-stratified analyses.

**Supplemental Figure 2:** Expression of *MCF2*, *HDAC8*, *FTX*, and *SLC10A3* in males and females

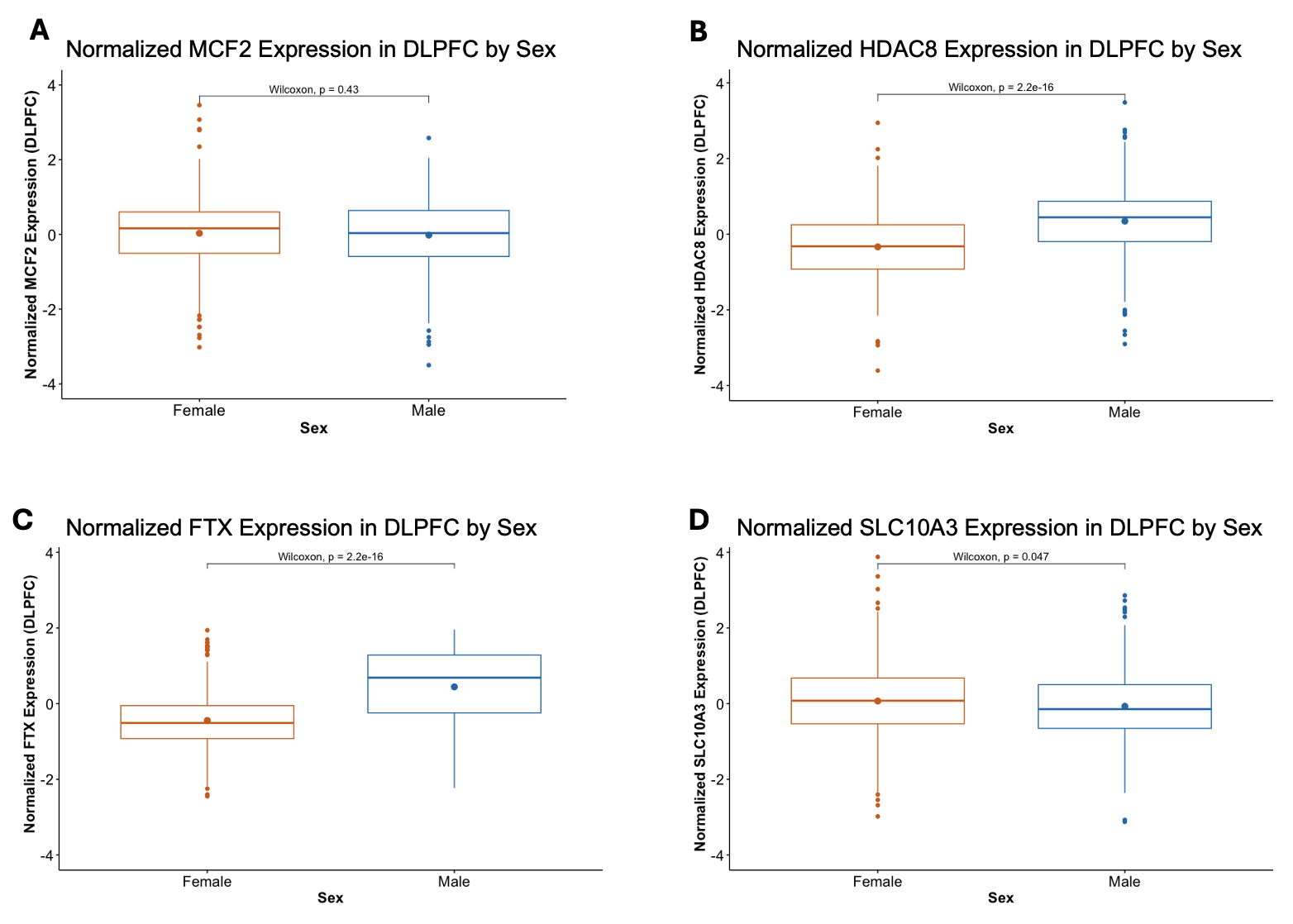

Box plots summarizing the normalized expression of *MCF2*, *HDAC8*, *FTX*, and *SLC10A3* in the DLPFC. Females are in orange and males are in blue. Expression was compared between males and females. Wilcoxon p-values are provided for each gene.

**Supplemental Figure 3:** Autosomal and X chromosome RNA sequencing quality control (QC) flow chart

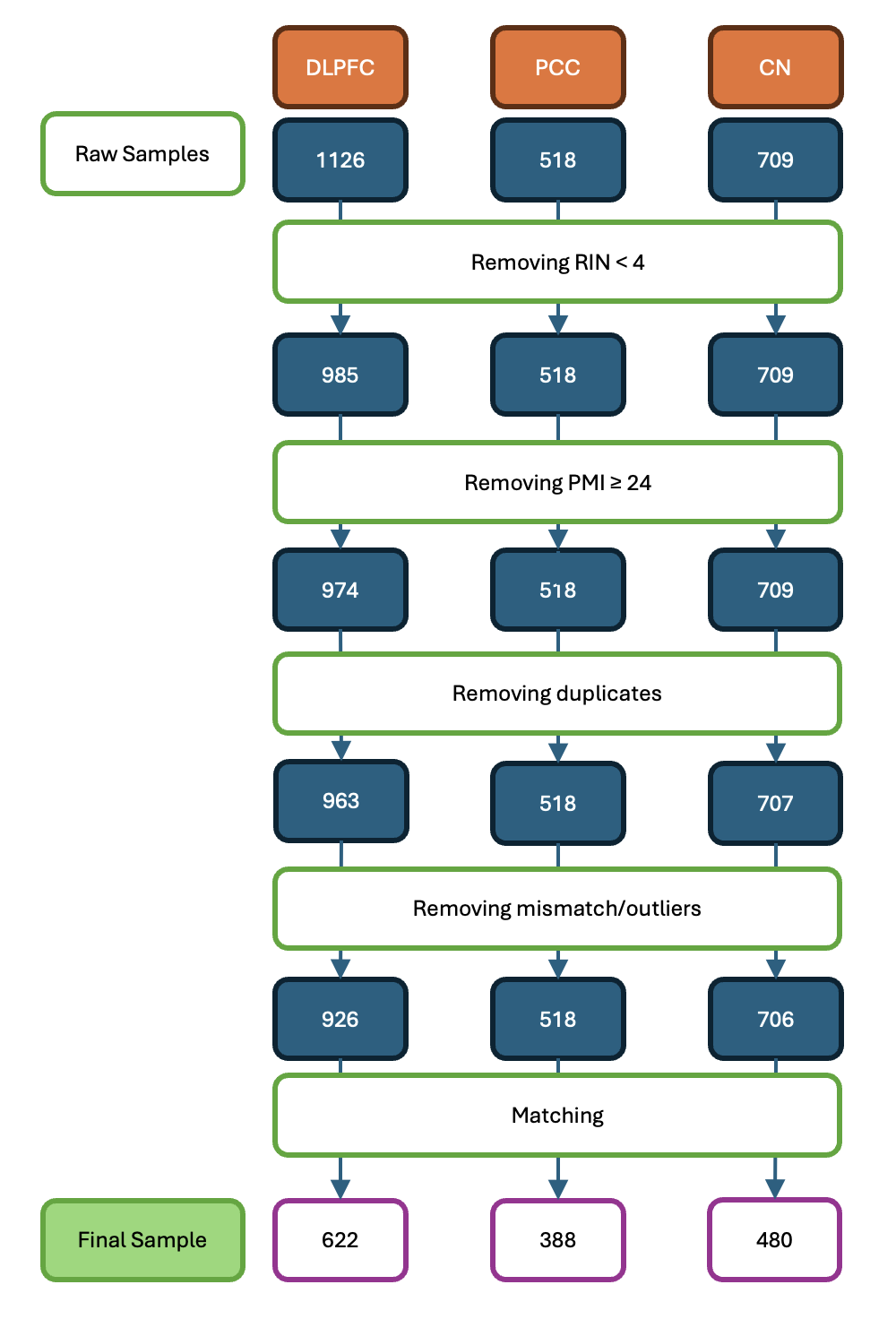

A flow chart depicting the number of participants remaining after each step of RNA sequencing quality control. The final, matched sample used for analysis is depicted at the bottom.

**Supplemental Figure 4:** Y chromosome RNAseq QC flow chart

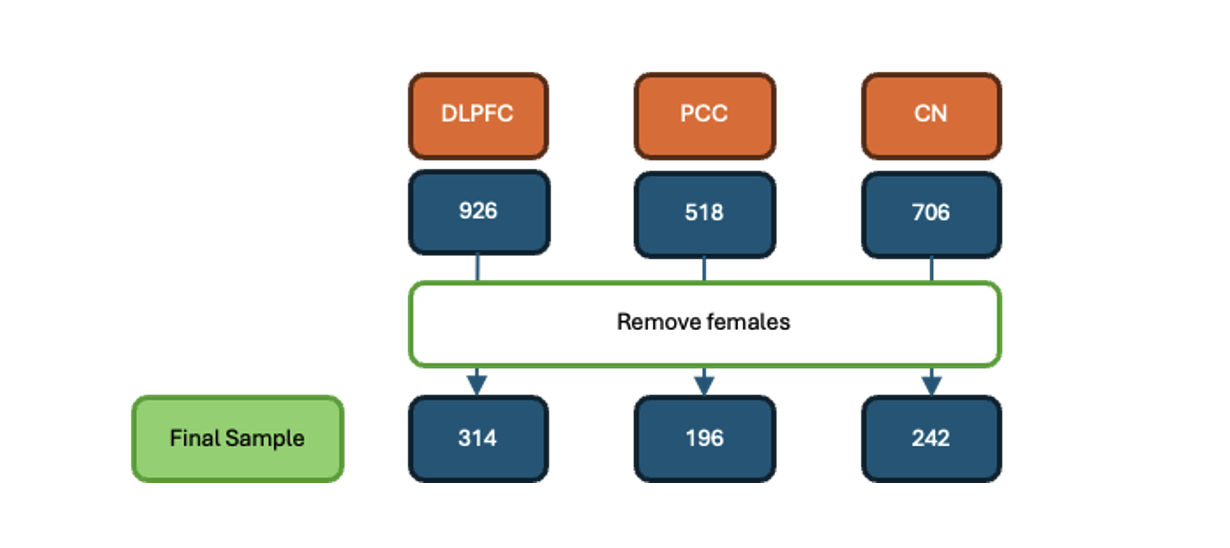

A flow chart depicting the number of participants remaining after each step of RNA sequencing quality control. Females were excluded as they do not have a Y chromosome. The final sample used for analysis is depicted at the bottom.

**Supplementary Table 1: Demographics for all participants**

|  | **TOTAL** |
| --- | --- |
| N | 767 |
| Age at Death (years) | 88.26±6.54 |
| Education (years) | 16.76±3.49 |
| Follow-up (years) | 7.41±4.92 |
| Latency to Death (years) | 0.74±0.95 |
| Post-mortem Interval (hours) | 7.62±4.45 |
| % Normal Cognition | 34 |
| % APOEε4 positive | 26 |
| % Non-Hispanic White | 98 |
| Amyloid (mm^2^) | 3.98±4.09 |
| Tau Tangles (mm^2^) | 6.28±8.00 |
| Cognition at Last Visit | -0.73±1.05 |
| Annual Change in Cognition | -0.11±0.10 |

**Supplementary Table 4: Summary table including sensitivity analyses for significant *sex x X-linked* gene interactions in the DLPFC**

| Outcome | Gene | Analysis | β | SE | T | P_Uncorrected_ |
| --- | --- | --- | --- | --- | --- | --- |
| Tau Tangles | *MCF2* | **Sex *x MCF2*** | **0.28** | **0.07** | **3.74** | **0.0002** |
|  |  | Sex *x MCF2* + *APOE*ε4 | 0.26 | 0.07 | 3.65 | 0.0003 |
|  |  | Sex *x MCF2* + Aβ | 0.19 | 0.07 | 2.94 | 0.003 |
|  |  | Male-stratified | 0.10 | 0.05 | 1.94 | 0.052 |
|  |  | Male-stratified + *APOE*ε4 | 0.08 | 0.05 | 1.74 | 0.08 |
|  |  | Male-stratified + Aβ | 0.07 | 0.04 | 1.60 | 0.11 |
|  |  | **Female-stratified** | **-0.18** | **0.06** | **-3.33** | **0.001** |
|  |  | Female-stratified + *APOE*ε4 | -0.18 | 0.05 | -3.29 | 0.001 |
|  |  | Female-stratified + Aβ | -0.12 | 0.05 | -2.42 | 0.02 |
|  | *HDAC8* | ***Sex x HDAC8*** | **-0.30** | **0.08** | **-3.82** | **0.0001** |
|  |  | Sex *x HDAC8* + *APOE*ε4 | -0.29 | 0.088 | -3.86 | 0.0001 |
|  |  | Sex *x HDAC8* + Aβ | -0.20 | 0.07 | -2.87 | 0.004 |
|  |  | Male-stratified | -0.08 | 0.05 | -1.60 | 0.11 |
|  |  | Male-stratified + *APOE*ε4 | -0.08 | 0.05 | -1.61 | 0.11 |
|  |  | Male-stratified + Aβ | -0.03 | 0.05 | -0.65 | 0.51 |
|  |  | **Female-stratified** | **0.22** | **0.06** | **3.75** | **0.0002** |
|  |  | Female-stratified + *APOE*ε4 | 0.21 | 0.05 | 3.74 | 0.0002 |
|  |  | Female-stratified + Aβ | 0.17 | 0.05 | 3.39 | 0.001 |
| Longitudinal Cognition | *SLC10A3* | ***Sex x SLC10A3*** | **0.04** | **0.01** | **3.69** | **0.0002** |
|  |  | Sex *x SLC10A3* + *APOE*ε4 | 0.04 | 0.01 | 3.69 | 0.0002 |
|  |  | Sex *x SLC10A3* + Aβ + tau | 0.04 | 0.01 | 3.62 | 0.0003 |
|  |  | Male-stratified | 0.02 | 0.01 | 2.51 | 0.01^a^ |
|  |  | Male-stratified + *APOE*ε4 | 0.02 | 0.01 | 2.48 | 0.01 |
|  |  | Male-stratified + Aβ + tau | **0.02** | 0.01 | 2.31 | 0.02 |
|  |  | **Female-stratified** | **-0.02** | **0.01** | **-2.7.** | **0.006** |
|  |  | Female-stratified + *APOE*ε4 | -0.02 | 0.01 | -2.78 | 0.01 |
|  |  | Female-stratified + Aβ + tau | -0.02 | 0.01 | -2.84 | 0.005 |
|  | *FTX* | ***Sex x FTX*** | **0.04** | **0.01** | **3.82** | **0.0001** |
|  |  | Sex *x FTX* + *APOE*ε4 | 0.04 | 0.01 | 3.85 | 0.0001 |
|  |  | Sex *x FTX* + Aβ + tau | 0.04 | 0.01 | 3.31 | 0.001 |
|  |  | **Male-stratified** | **0.02** | **0.01** | **3.58** | **0.0004** |
|  |  | Male-stratified + *APOE*ε4 | 0.02 | 0.01 | 3.53 | 0.0004 |
|  |  | Male-stratified + Aβ + tau | 0.03 | 0.01 | 3.70 | 0.0002 |
|  |  | Female-stratified | -0.02 | 0.01 | -2.06 | 0.04 |
|  |  | Female-stratified + *APOE*ε4 | -0.02 | 0.01 | -2.22 | 0.03 |
|  |  | Female-stratified + Aβ + tau | -0.01 | 0.01 | -1.31 | 0.19 |

All presented p-values are uncorrected. Bolded values represent those that survived FDR-correction in discovery analyses.

P-values for sensitivity analyses are not corrected.

**Supplementary Table 5: Results for *sex x X-linked gene x APOE-ε4* interaction analyses**

| Outcome | Gene | Analysis | β | SE | T | P_uncorrected_ |
| --- | --- | --- | --- | --- | --- | --- |
| Tau Tangles | *MCF2* | Sex x *MCF2* x *APOE*ε4 | 0.03 | 0.16 | 0.18 | 0.86 |
|  | *HDAC8* | Sex *x HDAC8* *x APOE*ε4 | -0.25 | 0.18 | -1.40 | 0.16 |
| Longitudinal Cognition | *SLC10A3* | Sex *x SLC10A3 x APOE*ε4 | -0.04 | 0.02 | -1.75 | 0.08 |
|  | *FTX* | **Sex *x FTX* *x APOE*ε4** | **0.05** | **0.02** | **2.20** | **0.03** |
|  |  | *APOE*ε4-negative females | -0.01 | 0.01 | -1.48 | 0.14 |
|  |  | *APOE*ε4-positive females | -0.04 | 0.02 | -1.66 | 0.10 |
|  |  | ***APOE*ε4-negative males** | **0.01** | **0.01** | **1.96** | **0.0498** |
|  |  | ***APOE*ε4-positive males** | **0.05** | **0.02** | **2.85** | **0.01** |
